# Quantitative image analysis pipeline for detecting circulating hybrid cells in immunofluorescence images with human-level accuracy

**DOI:** 10.1101/2023.08.24.554733

**Authors:** Robert T. Heussner, Riley M. Whalen, Ashley Anderson, Heather Theison, Joseph Baik, Summer Gibbs, Melissa H. Wong, Young Hwan Chang

## Abstract

Circulating hybrid cells (CHCs) are a newly discovered, tumor-derived cell population identified in the peripheral blood of cancer patients and are thought to contribute to tumor metastasis. However, identifying CHCs by immunofluorescence (IF) imaging of patient peripheral blood mononuclear cells (PBMCs) is a time-consuming and subjective process that currently relies on manual annotation by laboratory technicians. Additionally, while IF is relatively easy to apply to tissue sections, its application on PBMC smears presents challenges due to the presence of biological and technical artifacts. To address these challenges, we present a robust image analysis pipeline to automate the detection and analyses of CHCs in IF images. The pipeline incorporates quality control to optimize specimen preparation protocols and remove unwanted artifacts, leverages a β-variational autoencoder (VAE) to learn meaningful latent representations of single-cell images and employs a support vector machine (SVM) classifier to achieve human-level CHC detection. We created a rigorously labeled IF CHC dataset including 9 patients and 2 disease sites with the assistance of 10 annotators to evaluate the pipeline. We examined annotator variation and bias in CHC detection and then provided guidelines to optimize the accuracy of CHC annotation. We found that all annotators agreed on CHC identification for only 65% of the cells in the dataset and had a tendency to underestimate CHC counts for regions of interest (ROI) containing relatively large amounts of cells (>50,000) when using conventional enumeration methods. On the other hand, our proposed approach is unbiased to ROI size. The SVM classifier trained on the β-VAE encodings achieved an F1 score of 0.80, matching the average performance of annotators. Our pipeline enables researchers to explore the role of CHCs in cancer progression and assess their potential as a clinical biomarker for metastasis. Further, we demonstrate that the pipeline can identify discrete cellular phenotypes among PBMCs, highlighting its utility beyond CHCs.

## Introduction

Cancer is the second leading global cause of death, and despite notable advancements in treatment, significant challenges persist^1^. Timely information regarding disease burden is crucial for early detection, monitoring treatment response, and assessing the risk of metastasis or recurrence. However, current state-of-the-art assays and approaches for procuring such information pose difficulties. Biopsies and molecular analysis of tumor samples offer valuable insights but are invasive, limited in scope, and not feasible after tumor resection^2^. Alternatively, blood-based biomarkers hold promise, as they can be monitored from a peripheral blood sample, which is less invasive and facilitates more frequent monitoring during treatment. Current blood-based cancer biomarkers include non-cell-based biomarkers (i.e., cell-free nucleotides, proteins, exosomes) and cell-based biomarkers^3^. Non-cell-based cancer biomarkers, such as the protein, carcinoembryonic antigen (CEA), in colorectal cancer (CRC), are used for monitoring disease progression^4^, but are not always accurate or detected in every patient. Further, other biomarkers such as cell-free DNA (cfDNA) offer only single-dimensional information about the disease; for example, only small amounts of genomic information can be gained from cfDNA without additional tumor-associated protein information, and vice versa for protein-based biomarkers such as CEA^3^. Cell-based biomarkers are directly disseminated from the tumor and allow for a wider range of information to be assessed (i.e., RNA, genomic, and proteomic data)^3^. Circulating tumor cells (CTCs) are the most well-researched cell-based biomarker for cancer^5–9^. However, they are rare in peripheral blood (<5 CTCs/7.5 mL of blood in high burden settings), which makes them difficult to isolate and assess. Further, their numbers do not always correlate with patient outcomes^5–9^.

An exciting and emerging cell-based analyte, circulating hybrid cells (CHCs), is a newly discovered population of disseminated tumor cells that co-express immune and neoplastic proteins, and show promise as a cell-based biomarker^10,11,12^. CHCs can result from a fusion event between an immune cell and a tumor cell, granting it novel traits, such as enhanced migratory properties, and increased metastatic potential^10^. CHCs have been identified as the predominant tumor cell population circulating in peripheral blood^12–20^ across various types of cancers. Additionally, the number of CHCs in a cancer patient’s peripheral blood correlates with disease stage and patient outcomes, solidifying their potential as a useful cancer biomarker^10–12,21^. To optimize the utility of this discovery, the development of an accessible platform capable of identifying and analyzing CHCs is required. Such a platform and subsequent analytics will provide the foundation to move CHC analyses from discovery research-based to clinically approved assays. Such assays would improve cancer management by providing valuable and actionable information for personalized and timely treatment decisions^5^.

Several FDA-approved and commercial platforms are available for the detection of CTCs, such as CellSearch^Ⓡ22^, Vortex^23^, and DEPArray^24^. CellSearch^Ⓡ^ utilizes immunomagnetic isolation, Vortex employs size-based microfluidic separation, and DEPArray utilizes dielectrophoresis to trap cells. However, these methods principally focus on CTC detection, limiting their ability to identify CHCs^12,25^ by excluding all cells expressing immune surface antigens. Immunofluorescence (IF) imaging is predominantly used as a CHC detection method in the laboratory setting and relies on labeling of peripheral blood mononuclear cells (PBMCs) with panels of antibodies against cancer- and immune-associated epitopes. IF imaging provides a cost-effective approach for detecting CHCs, and advanced multiplex imaging techniques like cyclic IF (cyCIF)^26^ allow for the measurement of >20 biomarkers, as compared to 3-5 biomarkers in traditional IF, enabling detailed phenotyping of cells^12,27^.

Current CHC detection approaches in IF images have various challenges. First, machine learning (ML)-based approaches struggle with the imbalanced cellular dataset^28^, as CHCs are relatively rare and typically constitute less than 1% of PBMCs as shown in **Figure 1A**. This limitation particularly affects the discovery of new phenotypes, and makes generating labeled data challenging. Second, IF images contain autofluorescence, variable intensity levels across samples or batches, and artifacts that heavily impact downstream analyses, necessitating robust quality control and normalization steps during image processing^29,30^. Moreover, a standardized IF imaging protocol for serum samples is yet to be established; few studies address batch variation in serum samples, therefore the lower dynamic range of biomarkers compared to tissue complicates computational thresholding methods^31^. The traditional approach to CHC detection in IF images involves manual gating of biomarker expression with microscopy software such as ZEN or ImageJ^32^. However, this approach is not only time-consuming, but the extent of inter-observer and intra-observer bias in CHC detection remains unknown (**Figure 1B**). One study with 11 participants reported an initial inter-observer agreement of 85%, refined to 93% with further training^33^. However, this study focused on CTC detection, which are defined as CD45^-^/PanCK^+^, as opposed to CHCs, which are defined as CD45^+^/PanCK^+^. Although current software solutions strive for fully automated CTC detection in IF images, some of them require specialized tools like functionalized and structured medical wire^34^ for CTC collection, making them unsuitable for CHCs and for users lacking access to such technologies. Few software offer solutions for CHC detection. Support vector machine (SVM) classifier ALICE can enumerate 20 CTC phenotypes, including CHCs^35^. However, ALICE’s threshold-based segmentation is not robust against known challenges in PBMC IF images, such as artifacts and cell clumps. Its performance also relies on first capturing potential CTCs with TU-chip^TM^, a microfluidic device for sorting cells by size, and has been tested for CHC detection on only pancreatic cancer datasets. Given that the size of CHCs is similar to leukocytes^10^, it is most likely that the TU-chip^TM^ is missing the majority of CHCs. Further, ALICE does not critically address human inter-observer and intra-observer bias in ground truth (GT) generation.

**Figure 1.**
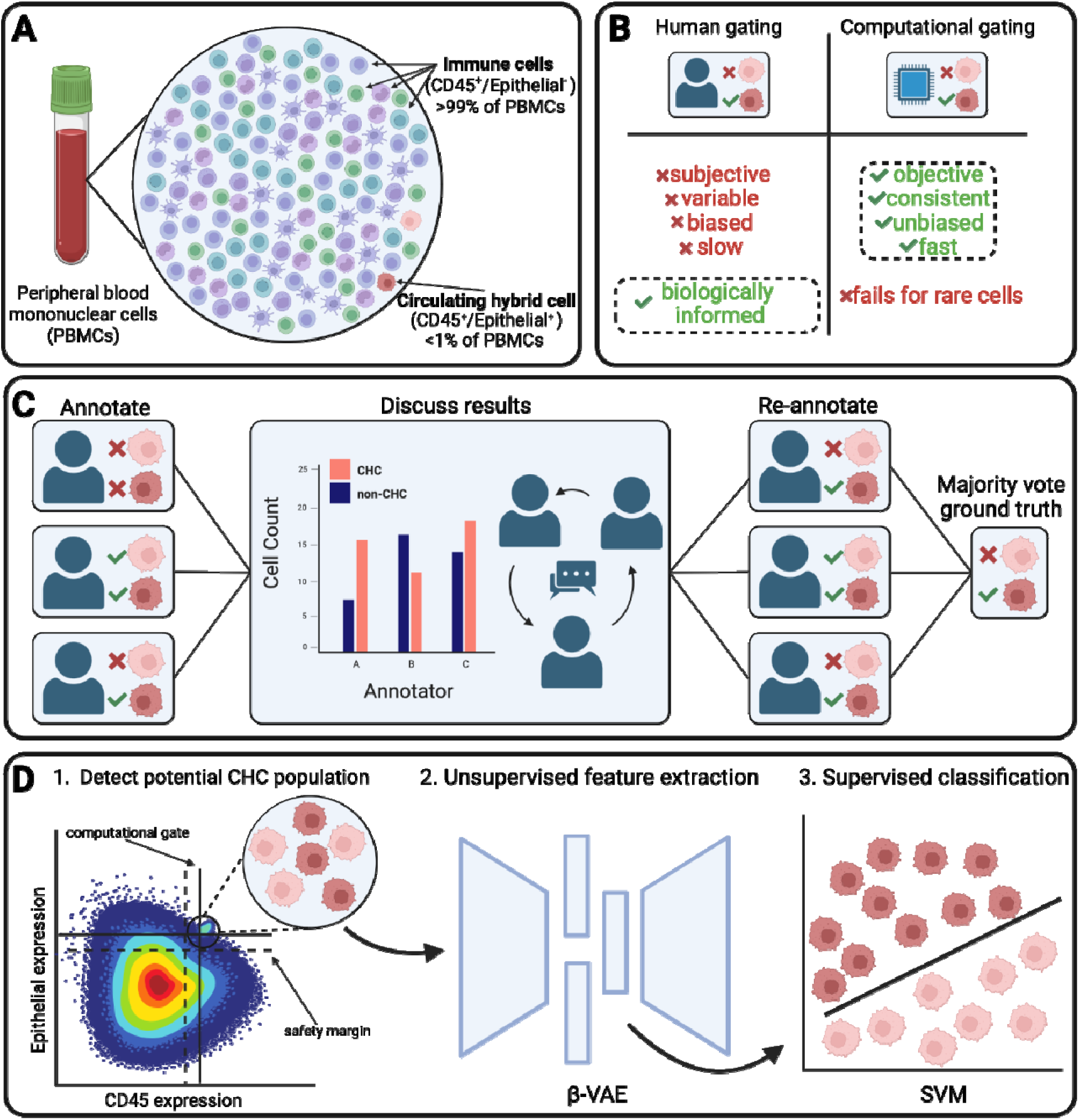
Pipeline motivation and overview. **(A)** CHCs are a rare cell population in peripheral blood. **(B)** The benefits and drawbacks of human and CG methods for inferring cell types in IF images. **(C)** Our rigorous, iterative human annotation study to determine GT labels. **(D)** Our image analysis pipeline that uses CG to filter potential CHCs to a β-VAE representation learner, which extracts latent features from the single-cell images to be classified with a support vector machine (SVM).

Herein, we introduce an automated CHC detection pipeline that achieves human-level performance. Our pipeline combines statistics-driven computational gating (CG) and representation learning to effectively identify and phenotype CHCs. We conducted a thorough human annotation study to establish accurate ground truth (GT) CHC labels. To manage the large initial dataset, encompassing 1 million PBMCs, comprising 9 patient samples from 2 disease sites (**Table S1**), we first employed CG to narrow the analytical dataset to 2,834 PBMCs for human annotators to review (**Figure 1C**). We provided the single-cell data in a format that aligns with the annotator’s familiar view of stained cells, including negative controls. We examined annotator variation and bias through a systematic evaluation of 7 annotator’s performance. The initial results were shared with the annotators, allowing them to learn from any gating mistakes and phenotype calling biases. Annotators attended a review of the data and developed guidelines for CHC calling. To establish a robust GT dataset, we then carried out a second round of individual annotations with the original 7 annotators and 3 additional annotators, all of who attended the initial data review (**Figure 1C**). We then use a β-VAE to extract latent features from the single-cell images and train an SVM classifier to detect the CHCs (**Figure 1D**). Finally, we compare the PBMC classification performance of CG alone, the SVM, and the human annotators.

## Results

### Image pre-processing and computational gating strategy

The primary obstacle to developing a robust model for CHC detection lies in obtaining a well-annotated dataset. Unfortunately, there is currently no publicly available PBMC dataset with adequate CHC annotations. To address this issue, we created our own labeled dataset, enlisting 10 individuals experienced in annotating CHCs in IF images. Our collected PBMC dataset consists of samples from 9 patients, featuring 15 regions of interest (ROIs) and a total of 1,083,196 segmented PBMCs. Annotating such a vast number of cells with the manual efforts of 10 people would have been impractical. To overcome this challenge, we implemented deep-learning based cell and nuclei segmentation, Mesmer^36^, feature extraction, and a CG strategy (**Figure 1D**) to filter the dataset through the expression pattern of an epithelial-directed antibody cocktail containing antibodies against E-cadherin (ECAD), epithelial cellular adhesion molecule (EpCAM), and pan-cytokeratin (panCK), as well as the pan-leukocyte antigen CD45, and their cellular morphology, allowing annotators to focus solely on potential CHCs. By applying this filtering approach, we significantly reduced the dataset to 2,834 PBMCs, accounting for only 0.26% of the original cell population. Each annotator assessed the samples on a per-ROI basis (**Methods**). Implementing this filtering strategy led to a remarkable reduction in annotation time, saving hours of manual effort and enabling annotators to concentrate more on evaluating the protein expression pattern within the potential CHCs.

### Annotation study

The need for a reliable labeled dataset gave us the opportunity to assess inter-annotator variation and annotator bias. We conducted two rounds of annotations to combat the effect of annotator disagreement. We chose majority vote aggregation (6 out of 10) from round 2 to define the GT labels (**Figure 1C**).

Annotator F1 scores showed a significant improvement between rounds 1 and 2, as seen in **Figure 2A**: For each ROI, median F1 scores increased and the interquartile range decreased. Annotators tended to be more conservative in identifying CHCs in round 2, as shown in **Figure S1**. **Figures S2** and **S3** further elucidate the inter-annotator variability in changes in F1 score and CHC count across annotation rounds. Our findings demonstrate that after human annotator training (**Methods**), annotators achieved a higher level of consensus on cell type assignments, increasing from 40.4% to 65.7% of the dataset, as shown in **Figure 2B**. This improvement highlights the significance of training in enhancing the consistency and accuracy of annotations.

**Figure 2.**
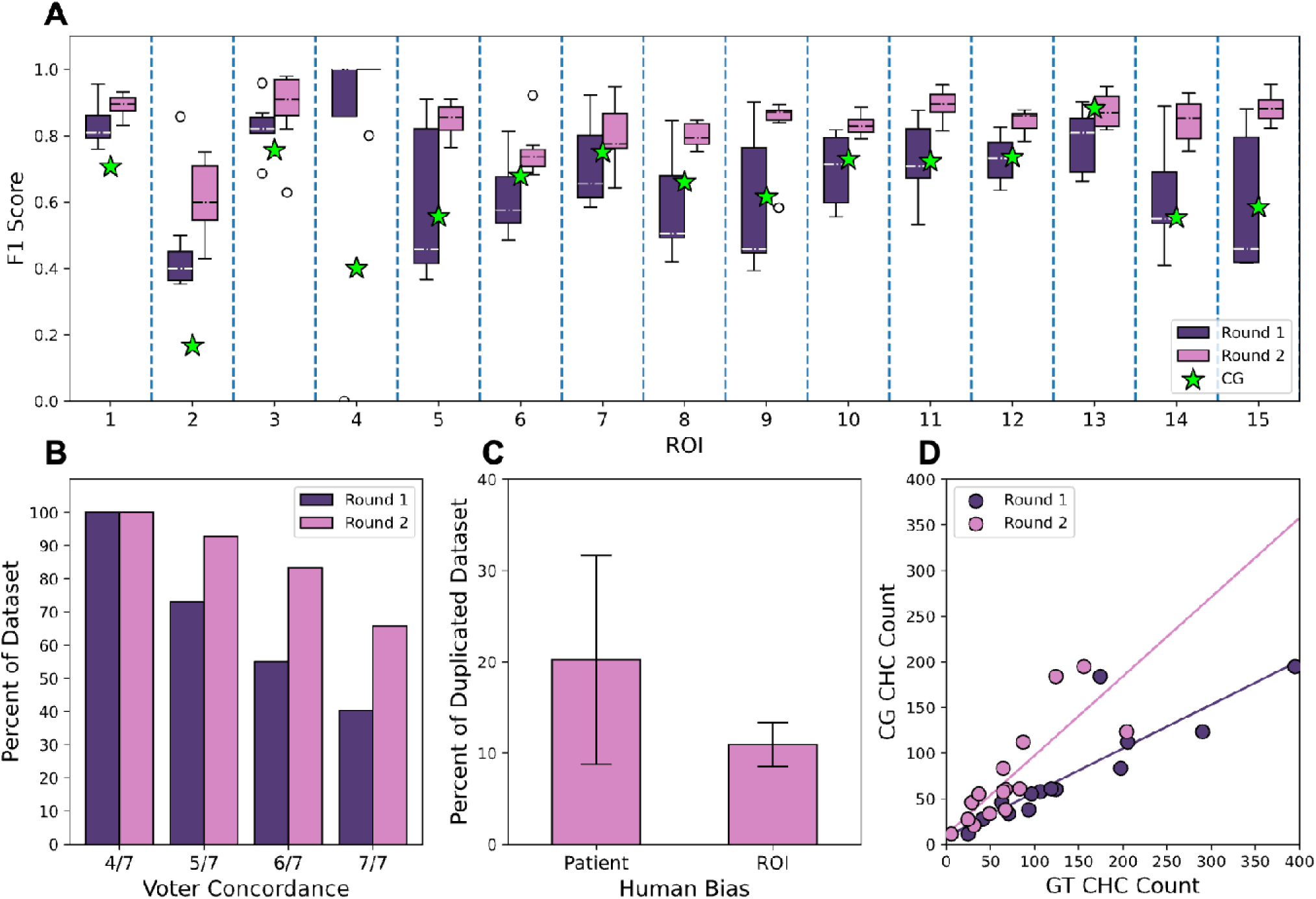
Human annotator variation, bias, and alignment with CG. **(A)** F1 score box-and-whisker plots for round 1 and round 2 per ROI. Circles mark outliers and the stars mark the CG score. **(B)** Inter-annotator agreement across rounds, where each bar is the fraction of cells in the filtered dataset that received greater than or equal to the number of votes. **(C)** Patient and scene bias of 7 annotators who completed round 1 based on the duplicated dataset (Methods), with error bars representing ±1 standard deviation. **(D)** CG CHC count plotted against the human-generated GT counts for rounds 1 and 2, with corresponding linear regression lines.

Next, we assessed annotator bias introduced from patient-level and ROI-level variations (**Figure 2C**). Initially, we observed that, on average, the annotator’s cell type assignments changed for approximately 20% of the cells duplicated across different patients (**Methods**). This outcome was expected and likely influenced by the batch effect present among patients with varying disease states and data acquisition factors such as processing time, staining, and imaging. Surprisingly, we also discovered that, on average, the annotator’s cell type assignments changed for about 11% of the cells duplicated across different ROIs within the same patient slide. This level of ROI bias presents significant challenges to the conventional control well-based gating approach, which involves defining IF channel (protein expression) contrast limits for multiple ROIs of the same patient slide, relying on a single, unstained ROI to estimate autofluorescence levels.

While single-cell accuracy holds significant importance to biological investigations and cell phenotyping, the clinical use of CHCs to infer disease state and trajectory primarily relies on CHC enumeration^10,11,21,37^. Consequently, we compared CG against the GT counts of both rounds to assess its performance in capturing the overall trend of CHC presence within the ROIs. We use the majority vote (4 out of 7) from round 1 as the GT labels for comparison purposes. The results give an R^2^ of 0.68 and a slope of 0.87 for round 2, as shown in **Figure 2D**, indicating a substantial alignment between CG and GT counts. However, CG had a better regression to round 1 GT with an R^2^ of 0.77, highlighting the necessity for more advanced classification methods, especially in light of the observed elevated F1 score during round 2.

### Representation learning

Having established the accurate GT labels, we moved to CHC classification and phenotyping (**Figure 1D**). In short, our CG method measures the background signal to estimate the true positive signal and identifies positive cells by gating the fraction of positive pixels in the cell area (**Methods**). While useful for finding potential-CHCs, this approach is limited to the cell’s intensity profile and does not take into account spatial localization of expression, texture, and cell morphology^38^. Consequently, we applied a β-variational autoencoder (β-VAE) to the single-cell images, which successfully learned single-cell representations that effectively differentiated CHCs from other PBMCs. We subsequently trained an ML classifier to improve CHC detection in PBMC samples over CG.

In the uniform manifold approximation and projection (UMAP) visualization (**Figure 3A** and **3B**), the CG labels (color-coded in **Figure 3A**) showed lower class separation, with more positive samples appearing in the dominantly negative region. However, the GT labels (color-coded in **Figure 3B**) demonstrate improved separation between classes in the projection. In the class density plots of the CG-colored UMAP (inset in **Figure 3A**), substantial overlap among the classes was evident. In contrast, when stratified by annotator vote (**Figure 3C**), the same density plots maintained the positive/negative spatial distribution while showing clearer separation between high-confidence non-CHCs (0 out of 10 votes) and high-confidence CHCs (10 out of 10 votes). This underscores the use of representation learning to achieve clear class separations and capture annotator consensus compared to the CG approach. While some level of patient-induced batch effect was expected, the CHC region generally exhibited a well-mixed patient distribution as shown in **Figure S4A**. Importantly, the observed CHC enrichment was driven by biologically meaningful features, such as epithelial cocktail and CD45 expression (**Figure S4B**).

**Figure 3.**
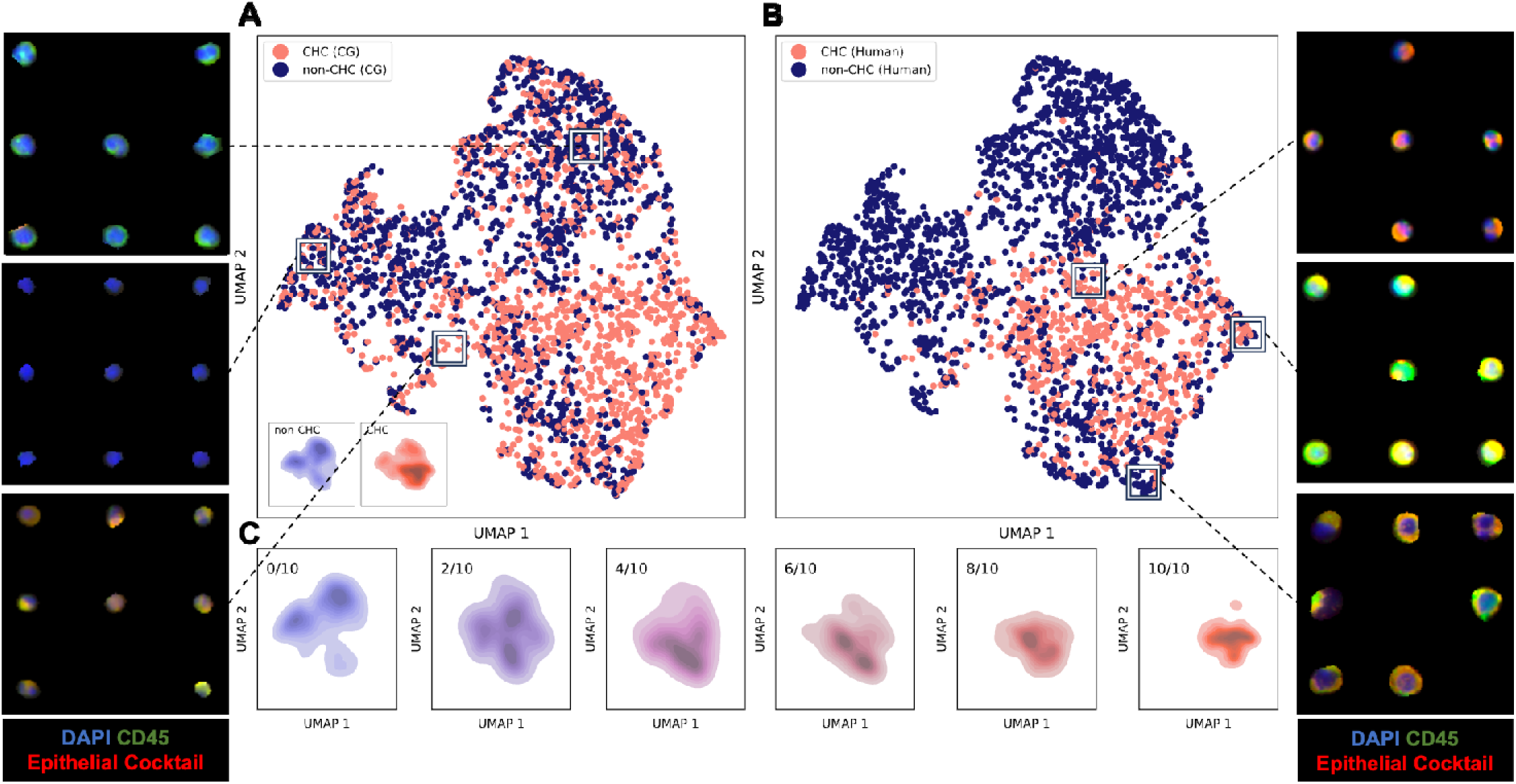
UMAP of single-cell. β**-VAE embeddings. (A)** UMAP of β-VAE embeddings from the filtered CHC dataset, colored by CG callouts with positive (salmon) and negative (midnight blue) density plots in the lower left corner. **(B)** UMAP of VAE embeddings from the filtered CHC dataset colored by the human GT callouts. **(C)** Density plots of annotator agreement, ranging from cells with 0 votes (confident negative) to 10 votes (confident positive). Additionally, single-cell images plotted in the UMAP space show clusters of morphologically distinct cells. To view the full single-cell image UMAP, visit https://heussner.github.io/pbmc-umap/.

Furthermore, the single-cell representations obtained from the β-VAE exhibit the ability to differentiate cell subtypes characterized by distinct morphologies and staining patterns. For example, six representative cell clusters are shown in **Figure 3A** (negative CHC region) and **3B** (positive CHC region), each displaying unique staining patterns and morphologies. The UMAP of all single-cell images can be seen in https://heussner.github.io/pbmc-umap/ for closer inspection. Notably, the β-VAE’s ability to learn morphological features enables it to identify irregularly shaped cells, which is particularly valuable for detecting poorly segmented cells.

### PBMC classification

Finally, we trained an SVM, a linear model, and a multilayer perceptron with single-cell embeddings to classify the PBMCs as CHCs or non-CHCs. We found that the SVM outperformed the other models at this task. We then compared its accuracy to the round 1 annotators, round 2 annotators, and CG, shown in **Table 1**. The SVM achieved near human-level accuracy on CHC detection, with a 5-fold cross-validation average F1-score of 0.80 with a top F1 score of 0.84. The round 2 annotators averaged an F1 score of 0.83 with a top F1 score of 0.84. A paired t-test on the 5-fold F1 scores revealed a t-statistic of 2.18 with a p-value of 0.095, finding no statistically significant difference between the human’s and the SVM’s cross-validation performance. Furthermore, the ROI-induced bias (11% of cell labels changed when the cell was moved to another ROI of the same patient sample) shown in **Figure 2C** provides additional support for the SVM’s ability to match human accuracy. The SVM also outperformed the untrained annotators and CG by a substantial margin.

**Table 1.** PBMC classification summary. PBMC classification performance summary, reporting average and top F1 scores of a 5-fold cross validation. Human F1 scores were averaged across the participating annotators for each fold, which were then averaged across the 5 folds. The CG and SVM F1 scores were averaged across the same 5 folds for a similar comparison.

| Method | Average F1 (SD) | Top F1 |
| --- | --- | --- |
| Human Round 1 | 0.60 (0.010) | 0.61 |
| Human Round 2 | 0.83 (0.0070) | 0.84 |
| CG | 0.61 (0.023) | 0.63 |
| SVM | 0.80 (0.027) | 0.84 |

## Discussion

Defining the GT labels for training ML models in biomedical applications is a well-known challenge^39^. While our labels are the product of 10 humans’ annotations over multiple iterations, we acknowledge that there are inherent limitations to our approach. Notably, the mosaic single-cell images (**Methods**) that were annotated differ from the typical images where cells sit next to each other on the slide that annotators usually work with, potentially introducing artificial biases into the annotation process.

To investigate this possibility, we acquired these conventional annotations for each ROI from a single annotator. We then compared these annotations with the GT counts and the CG counts (**Figure S5**). Our observations revealed a mixed pattern: Some ROIs exhibited higher conventional counts compared to the GT, while others showed lower counts. Upon further inspection, we find a strong negative correlation between the difference in conventional count and GT counts in relation to the size of ROI, as shown in **Figure 4**. Notably, ROIs containing over ∼50,000 PBMCs show lower conventional counts than GT counts. Conversely, ROIs with fewer than ∼50,000 PBMC ROIs demonstrated higher conventional counts relative to the GT. We posit that this trend underscores an ROI size-related bias inherent in the traditional cell enumeration approach, highlighting the potential oversight of certain CHCs populations during the labor-intensive inspection of individual cells within large ROIs.

**Figure 4.**
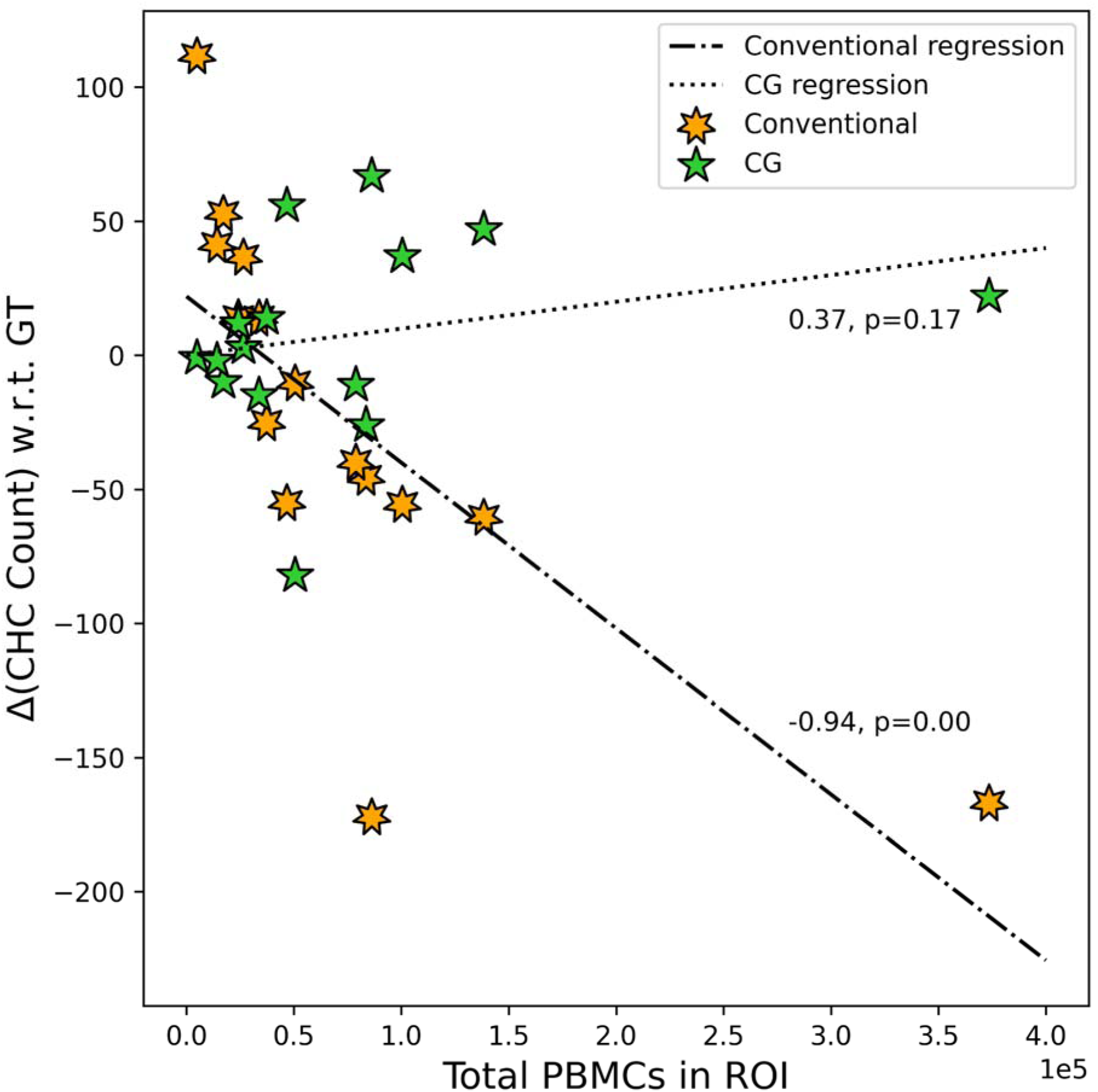
Conventional and CG CHC counts with respect to GT count versus ROI PBMC count. Differential CHC count of the conventional method and CG with respect to the GT counts, where +Δ is a higher count than GT and -Δ is a lower one. The dotted and dashed-dotted lines are linear regressions, and the text associated with each line is the Spearman’s rank correlation coefficient and p-value.

Another potential limitation of the study was the execution of the second round of annotations. In this round, a single annotator was responsible for setting contrast limits for 8 out of the remaining 9 annotators, thereby standardizing the epithelial cocktail and CD45 thresholds. However, it is noteworthy that annotator G deviated from these thresholds. Despite this, annotator G exhibited a comparable reduction in CHC count and an increase in F1 score between rounds 1 and 2, showing that the review session helped to mitigate variability regardless.

The amount of variation observed in CHC count across 7 annotators indicates the potential pitfalls of relying solely on a single annotator for CHC identification. Such reliance could lead to misleading results and potentially affect downstream analyses that aimed at predicting patient outcome^10^. Understandably, the task of labeling data is expensive, and involving multiple annotators might not always be feasible. However, our work highlights that annotator discrepancies largely decrease after training.

The annotation review session allowed the annotators to engage in constructive discussions concerning the staining intensity and morphological characteristics of CHCs. Detailed in **Methods**, the qualitative criteria adopted for CHC identification encompass various aspects, including the presence of artifacts, shape factors such as eccentricity and size, and the fraction of positive staining signals. The development of these guidelines resulted in enhanced consistency in the GT annotations. Notably, annotators who initially deviated significantly, such as annotator D or annotator G (**Figure S3**), were able to bring their annotations into better alignment with those of their peers. The annotation study provided this harmonization in a high-throughput and reproducible manner. Therefore, we advocate for the incorporation of similar opportunities for training in future studies that rely on human annotators. This practice could significantly enhance the reliability and validity of annotations, ultimately contributing to more robust research outcomes and greater reliability for clinical readouts.

The image analysis pipeline also has technical limitations that need to be addressed in future work. First, immunofluorescent contaminants of relatively high intensity can be erroneously identified as cells by segmentation algorithms. These artifacts lack a corresponding nucleus, and a simple approach involves matching the nucleus mask to the cell mask to eliminate those instances. However, occasionally the artifacts overlap true cells. To address these cases, the image is binarized at the estimated positive threshold, and outlier areas (>99% of measured cell area) are set to the estimated background level (**Methods**). Substantial overlap with true cells can occur, in which case the cell cannot be recovered for downstream analysis. Exceptionally large cells, such as cancer-associated macrophage-like cells (CAMLs), can be similar in size to the artifacts and are filtered out with this quality control (QC) step. In this study, we focus on CHC detection and not on cells that displayed CAML features. Thus, future work should aim for a better balance between artifact management and biological sensitivity.

Additionally, segmentation errors resulted in irregularly shaped cell areas. Aside from size, we did not exclude cells based on their morphology, aiming to encompass a broad spectrum of biological phenotypes, resulting in the inclusion of these imposter cells. For example, **Figure S6A** shows an imposter cell in row 5, column 1 of the mosaic. For the biologically informed annotators, these imposters were easily distinguished and not marked as false positives. However, avoiding the cells in the annotation study would have made space for more potential CHCs, improving the GT labels. In the case of cells in close proximity to each other, it would be useful to implement signal correction to account for lateral spillover across cell masks^40^.

To assess the SVM’s generalizability to new patient samples, we performed a 9-fold cross-validation, split by patient (**Figure S7**). The resulting average F1 score of 0.74 was low compared to the previous 5-fold cross-validation. The SVM performed worst on patient P2 (ROIs 2-4) with a score of 0.54 and best on patient P6 (ROI 9) with a score of 0.88. However, achieving high accuracy on held-out patients while only training on 8 patients is unreasonable and expected to improve with a larger, more representative dataset.

This image analysis pipeline was designed to support multiplexed IF imaging, thereby empowering researchers to explore clinically relevant CHC subtypes by leveraging 20+ biomarkers in combination with morphological and staining pattern differentiation^41^. This approach offers a more comprehensive and nuanced understanding compared to measuring the mean signal intensity of biomarkers. Moreover, β-VAE’s capability to learn staining patterns proves especially beneficial for CAMLs, which typically exhibit punctate epithelial cocktail staining as well as expression of CD45^42^. However, in this study, which focuses on CHC identification, abnormally large cells were filtered out as a QC step.

In summary, we combine CG with representation learning to detect CHCs in IF images, achieving near human-level accuracy. We demonstrate the poor agreement between individual annotators and reveal significant levels of bias in the annotation process. These observations call for future IF-based cell phenotyping studies to address these concerns. Our detection method and GT generation strategy may be applicable to many rare cell types and is not exclusive to serum samples. Our pipeline enables CHC research in a high-throughput, repeatable manner.

## Data and Software Availability

All single-cell images used in the annotation study will be publicly released (https://zenodo.org/record/8270693). All software used in this manuscript is detailed in the article’s Methods section and its Supplemental Information. The associated scripts are freely available via GitHub as described at https://github.com/heussner/crc-process.

## Acknowledgements

We thank Luke Strgar and Eun Na Kim for contributing to the image preprocessing and quality control code, Koei Chin for his advice on IF imaging, the Advanced Light Microscopy Core at OHSU, and Nicole Giske, Ranish Patel, John Swain, Abby Gillingham, Ethan Lu, and Ashvin Nair for help in annotating CHCs. This work was supported by the National Institutes of Health (R01 CA253860). YHC acknowledges funding from the National Institute of Health (U2CCA233280) and Kuni Foundation Imagination Grants, ANA acknowledges funding from the National Cancer Institute (F31CA271676). The resources of the Exacloud high-performance computing environment developed jointly by OHSU and Intel and the technical support of the OHSU Advanced Computing Center is gratefully acknowledged.

## Methods

### Human samples

All human specimens studies were approved by the Oregon Health & Science University (OHSU) Institutional Review Board under IRB protocol 5169. Informed consent was obtained from all subjects under IRB protocols for the Oregon Colorectal Cancer Registry, the OHSU Biolibrary, and the Oregon Pancreatic Tumor Registry. Peripheral blood samples were obtained from patients treated at OHSU (**Table S1**).

### Sample preparation, staining, and imaging

Patient peripheral blood was collected in heparin-coated vacutainer tubes and processed within 2 hours using standard approaches for PBMC isolation^11,12^. Briefly, whole blood was diluted with phosphate buffered saline (PBS) then subjected to density centrifugation with Ficoll-Paque PLUS (GE Healthcare, 17-1440-03) as previously described^21^. After isolation, PBMCs were spread onto glass slides coated with Poly-D Lysine (1 mg/mL, Millipore Sigma, A-003-E) at a density of 600,000 cells/slide, and allowed to dry for 15-30 min. Cells were then fixed for 20 min with 4% paraformaldehyde, washed with PBS 3 x 2 min, and subjected to increasing ethanol baths 70%, 90%, and 100% ethanol. Cell slides were then air-dried at room temperature in a dark compartment, and stored at 4°C. Antibody staining was performed by rehydrating cell slides, then blocking non-specific epitopes with BlockAid™ Blocking Solution (ThermoFisher). PBMCs were incubated with fluorescent-conjugated antibodies against ECAD, EpCAM, and panCK were diluted in BlockAid at 22°C for 45 mins or at 4°C for 16 h.Cells were counterstained with DAPI and coverslipped using fluoromount G (Invitrogen, 00-4958-02). Stained specimens were digitally scanned at 20x magnification with a ZEISS AxioScanner. Z1 with a Colibri 7 light source (Zeiss) equipped with the following filter cubes DAPI (Semrock, LED-DAPI-A-000), AF488 (Zeiss 38 HE), AF555 (Zeiss 43 HE), AF647 (Zeiss 50), and AF750 (Chroma 49007 ET Cy7). Regions of interest (ROIs) that avoided scanning and staining artifacts were selected and exported using ZEISS ZEN blue software version 3.4 (Carl ZEISS AG, Oberkochen, Germany).

### Image preprocessing

We utilized MCMICRO^43^ to stitch (ASHLAR)^44^ and correct background illumination (BaSiC)^45^. Next, we employed DeepCell’s Mesmer^36^ to segment the nuclei with the DAPI channel and cells with the projection of epithelial cocktail and CD45 markers.

### Quality control

One unforeseen challenge encountered with serum samples was the presence of irregular cell size distributions (**Figure S8 A**). To address this, we first developed a visualization tool to generate heatmaps plotting DAPI intensity, cell major axis length, and cell area on the original slide image (**Figure S8 B**). These heatmaps allow us to easily evaluate the adequacy of the blood smear protocol, staining process and image acquisition. Additionally, to remove imaging artifacts (**Figure S8C**), we filter out cells above the 99th percentile and below the 1st percentile in terms of cell size (**Figure S8 D**). We also excluded cells lacking corresponding nuclei to ensure data integrity.

### Computational gating

We gated epithelial cocktail and CD45 by first estimating their background levels by measuring the mean (L) and standard deviation (σ) of their corresponding image channel’s background intensity. Then, we set a positive threshold by adding 3σ to L. We multiplied the epithelial cocktail threshold by an adjustment factor of 1.5 to account for a low signal-to-noise ratio relative to CD45. Next, we compute the positive pixel ratio (PPR) for each cell, which represents the fraction of positive pixels to total pixels within the cell area. The PPR values were then thresholded using an elbow plot optimization approach. We confirmed the reproducibility of this strategy by comparing elbow curves for each ROI, as shown in **Figure S9**. These PPR thresholds were crucial in determining whether a cell was positive or negative for a given marker. While this step is tailored to CD45 and epithelial cocktail, it exhibited broad applicability and could be extended to other biomarkers for serum samples.

### Annotation study design

To carry out the annotation study, we curated a single-cell mosaic image containing *N* CG epithelial cocktail^+^/CD45^+^ cells from the previous step and the next *N* cells below the PPR threshold of epithelial cocktail^+^ (“potential-CHCs”) for each ROI. We chose epithelial cocktail as the limiting biomarker for selecting potential CHCs due to its relative rarity in positive expression compared to CD45; we expected >95% of PBMCs to be CD45^+^, while <1% of PBMCs to be epithelial cocktail^+^. We also added *N* negative control cells which were determined by an epithelial cocktail PPR of 0 (many cells) and CD45 PPR below the previously determined positive threshold (few cells). Negative controls comprised the top row of the mosaic, which the annotators used to set their histogram thresholds. Finally, to study annotator bias, we created a bank of CG CHCs and duplicated them into other ROI mosaics. Specifically, we duplicated cells and placed them in neighboring ROIs from the same patient sample (98 cells), and placed others in ROIs from different patient samples (746 cells). These comprised 20% of the total cells in each mosaic. We added cells to the mosaics by cropping them from the processed ROI and padding them to be 96 by 96 pixels. Each mosaic was a 16-bit RGBA image; epithelial cocktail (red), CD45 (green), DAPI (blue), and gridlines (alpha). We trained annotators to use ImageJ^32^ software to annotate the PBMCs as CHCs or non-CHCs with the paint tool. An example mosaic side-by-side with human annotations is shown in **Figure S6**.

### Histogram settings for CHC annotation

Visualization settings were adjusted for each ROI based on negative control cells. Specifically, the minimum and maximum values were set to just below what could be visibly detected in the negative control cells along the top row of the mosaic for both CD45 and epithelial cocktail channels. To account for differences in CHC calling due to histogram setting variation between annotators, the minimum and maximum values were set by a single annotator and distributed to other annotators prior to their annotation in round 2. Annotator G did not follow this protocol, instead setting their own histogram values.

### CHC annotation consensus

After completing the first round of CHC annotations, all annotators met to review the data and determine a set of consensus features for CHC calling. The set of criteria that was agreed upon include the following: 1) cells must have an intact nucleus with adequate DAPI signal (**Figure S10 A**); 2) signal distribution of CD45 and epithelial cocktail should be around the majority of the cell membrane/cytoplasm (i.e. cells with bright specks of signal should not be called) (**Figure S10 B**); 3) cells with very dim and diffuse CD45 or epithelial cocktail signal should not be called (**Figure S10 C**). The criteria were distributed to 3 additional annotators prior to completing round 2.

### CHC enumeration

All reported CHC counts were normalized to 50,000 PBMCs to fairly compare ROIs with varying image sizes and cell densities^10^.

### β-VAE

We trained the β-VAE^46^ on the CG potential-CHCs, excluding the negative control cells and cell duplicates used in the annotation study for a total of 2834 cells. Our β-VAE had a convolutional encoder with 5 layers of 32, 64, 128, 256, and 512 filters and mirrored decoder, with a latent dimension of 64. The 16-bit single-cell images were resized to 128 by 128 pixels and mapped to an intensity range of [0,1]. We employed a β scheduler during training, starting at β=0.001 and increasing in intervals of 0.001 every batch until β=2 was reached and remained at until training was complete^47^. The batch size was 4 and learning rate was 0.0005, with early stopping and a maximum number of epochs of 1000. The model was built with the PyTorch Lightning platform and trained on 1 NVIDIA A100 GPU.

### Support Vector Machine (SVM) classification

We employed an SVM as the PBMC classifier in our study. The SVM is a powerful supervised ML algorithm widely used for classification tasks and is provided in the Scikit-Learn Python library. For our implementation, we chose the polynomial kernel with a degree of 2, allowing for a quadratic relationship between features. The coefficient (coef0) was set to 1, providing a bias term for the polynomial kernel function. The regularization parameter (C value) was set to 1, controlling the trade-off between maximizing the margin and minimizing the classification error. Additionally, we set the gamma parameter to 1, influencing the reach of the SVM’s influence on the decision boundary. To assess the performance of our SVM model reliably, we conducted k-fold cross-validation where k was equal to 5. This approach partitions the dataset into five subsets, using four of them for training and one for testing in each iteration. The process is repeated five times, and the final evaluation was obtained by averaging the results, providing a robust estimate of the model’s generalizability. By employing the SVM with the specified parameters, we achieved the highest F1 score of all the models evaluated during the study.

## Supplemental Tables and Figures

**Table S1:**
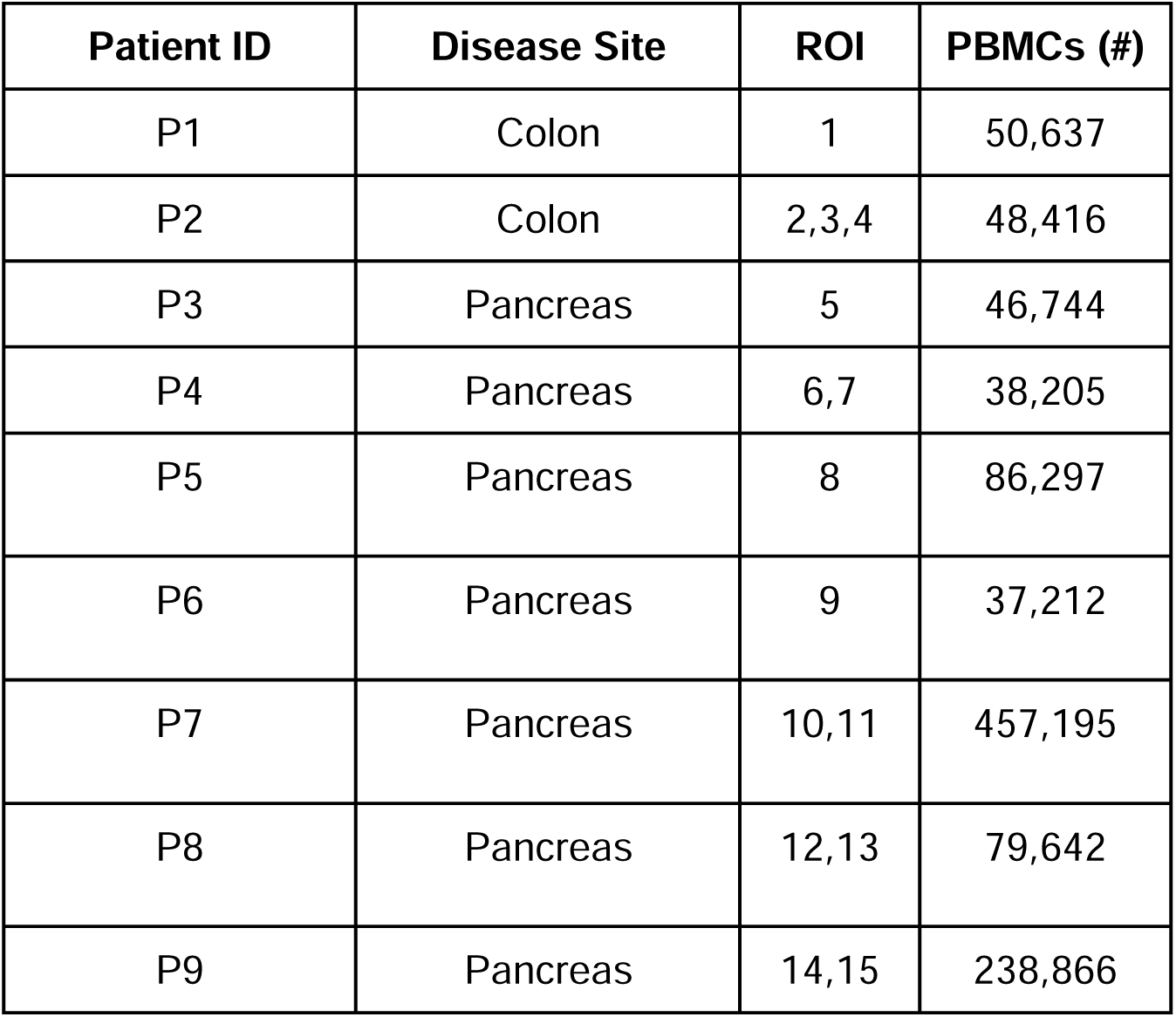
Dataset summary.

**Figure S1:**
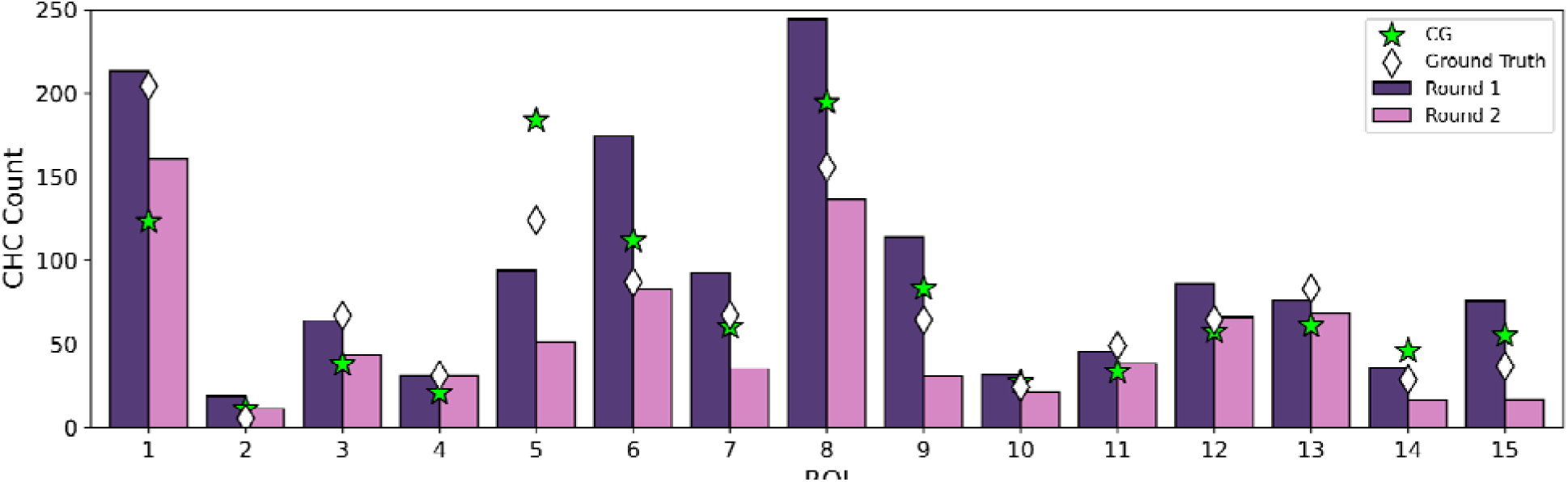
CHC counts per ROI across annotation rounds. Bars represent the median of 7 annotators who participated in both rounds. The diamond marks the majority vote GT and the star marks the CG count.

**Figure S2:**
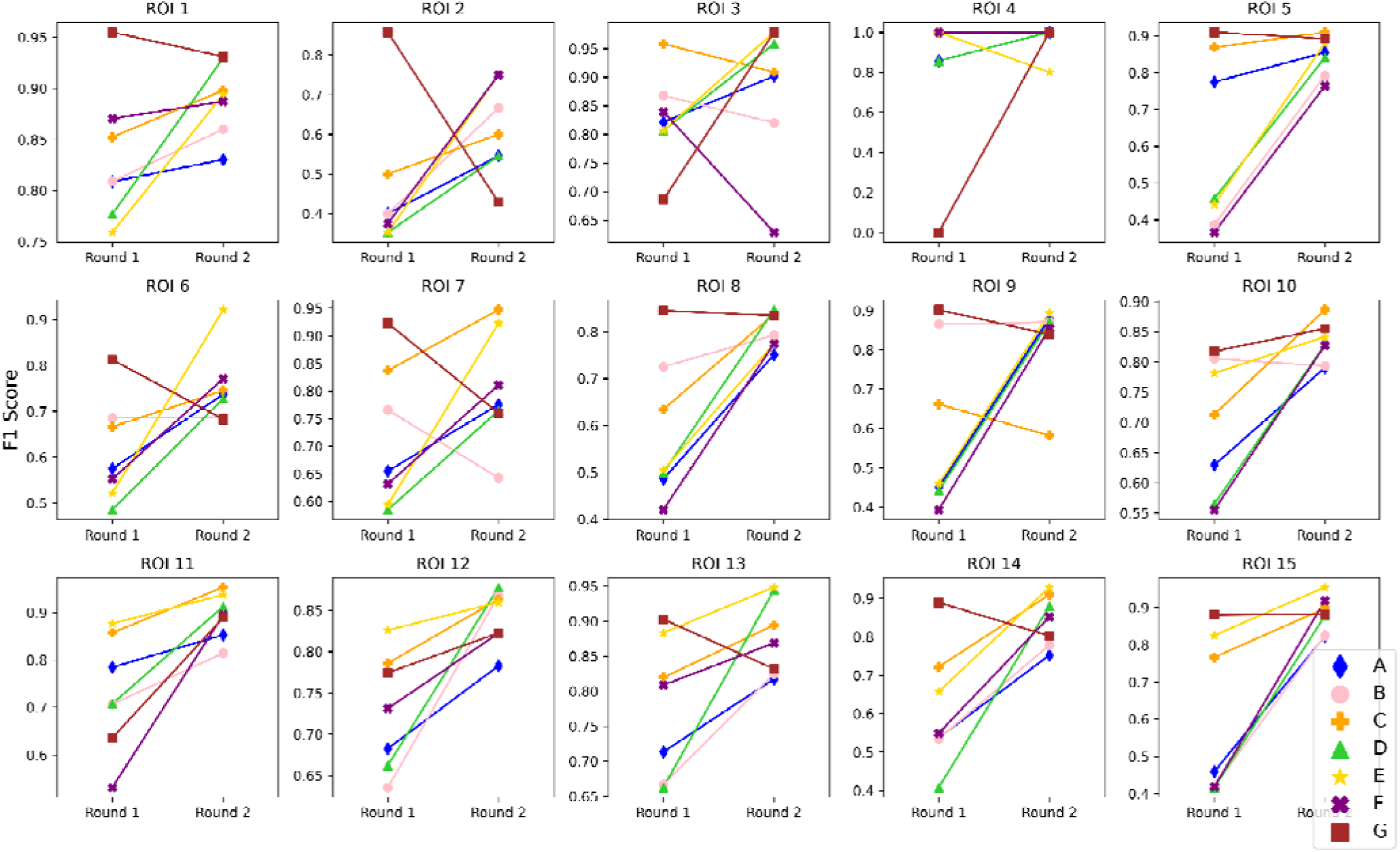
Annotator F1 score change across rounds for each ROI in the dataset.

**Figure S3:**
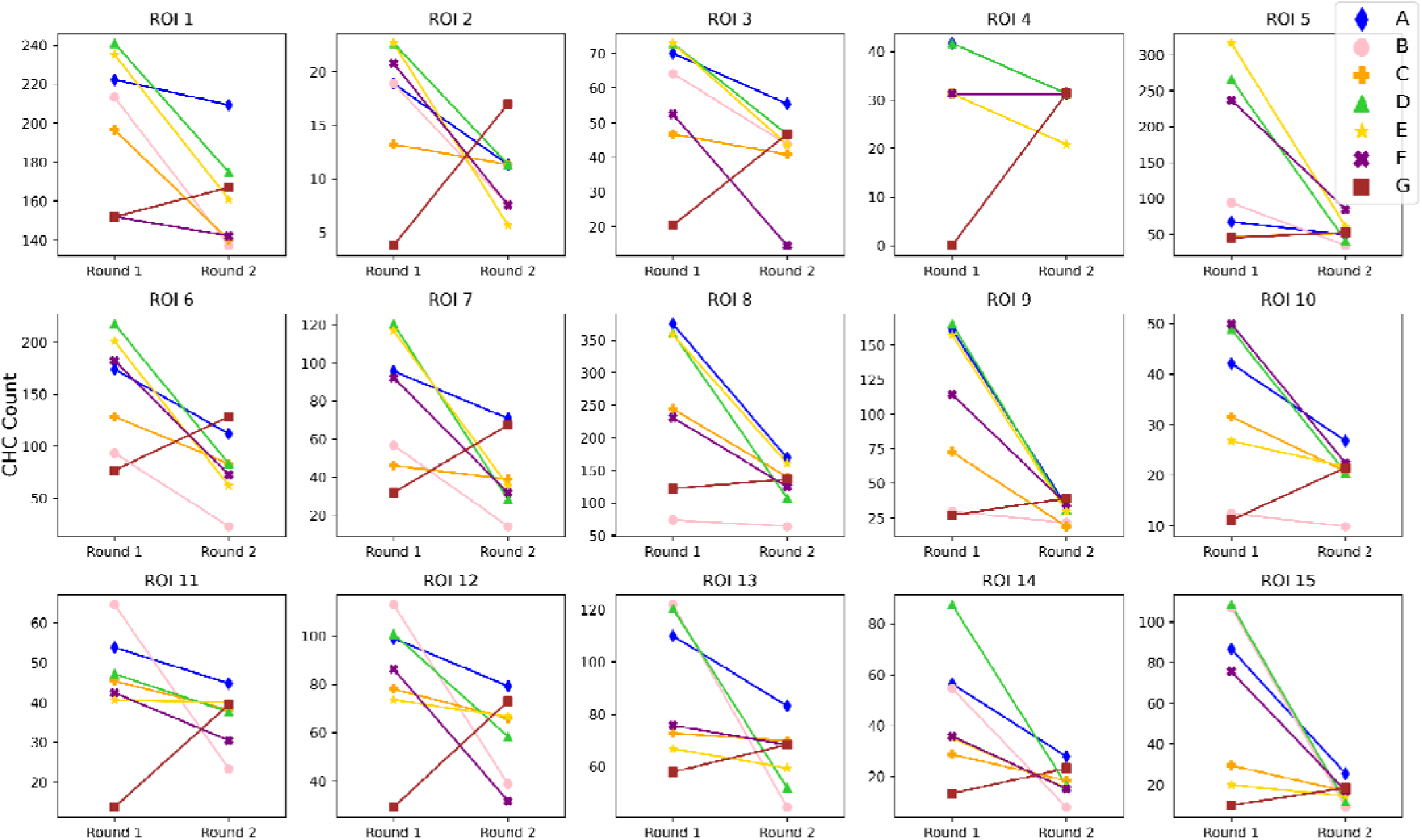
Annotator CHC count change across rounds for each ROI in the dataset.

**Figure S4:**
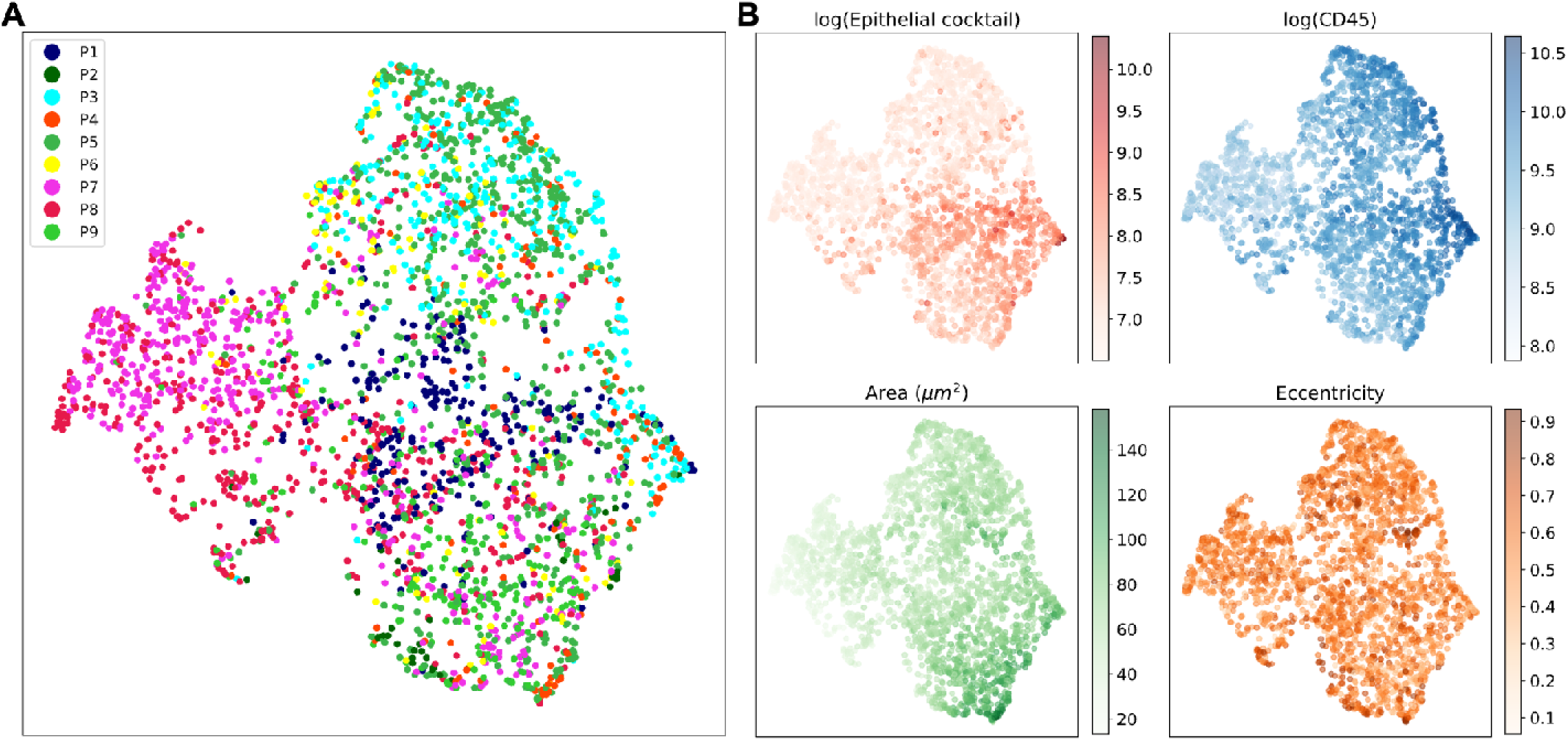
UMAP colored by patient and select features. **(A)** UMAP of single-cell VAE embeddings colored by patient. **(B)** UMAP of single-cell VAE embeddings colored by log-scaled epithelial cocktail mean intensity, log-scaled CD45 mean intensity, cell area in *um*^2^, and cell eccentricity.

**Figure S5:**
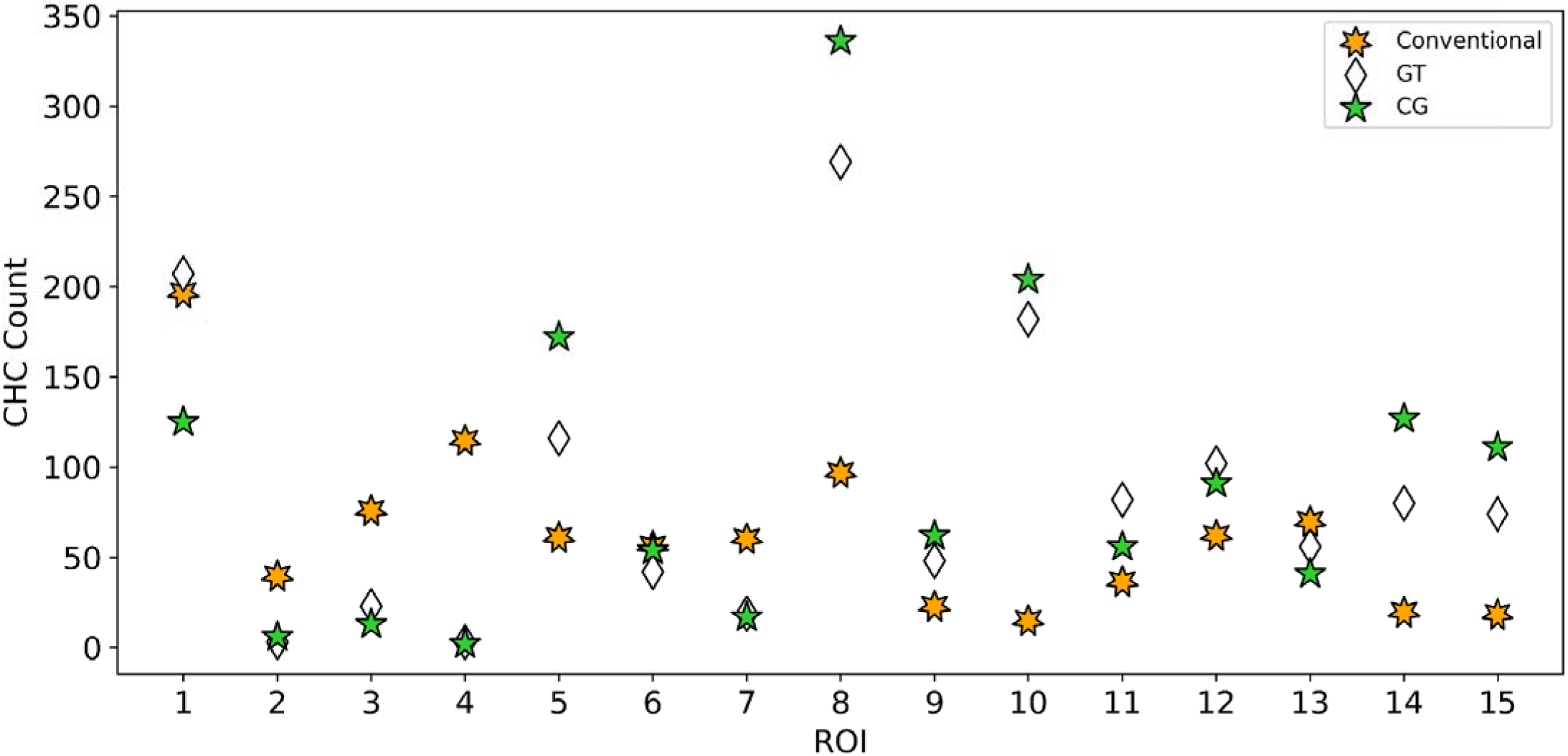
Conventional CHC counting method compared to the GT counts generated by the annotation study and CG.

**Figure S6:**
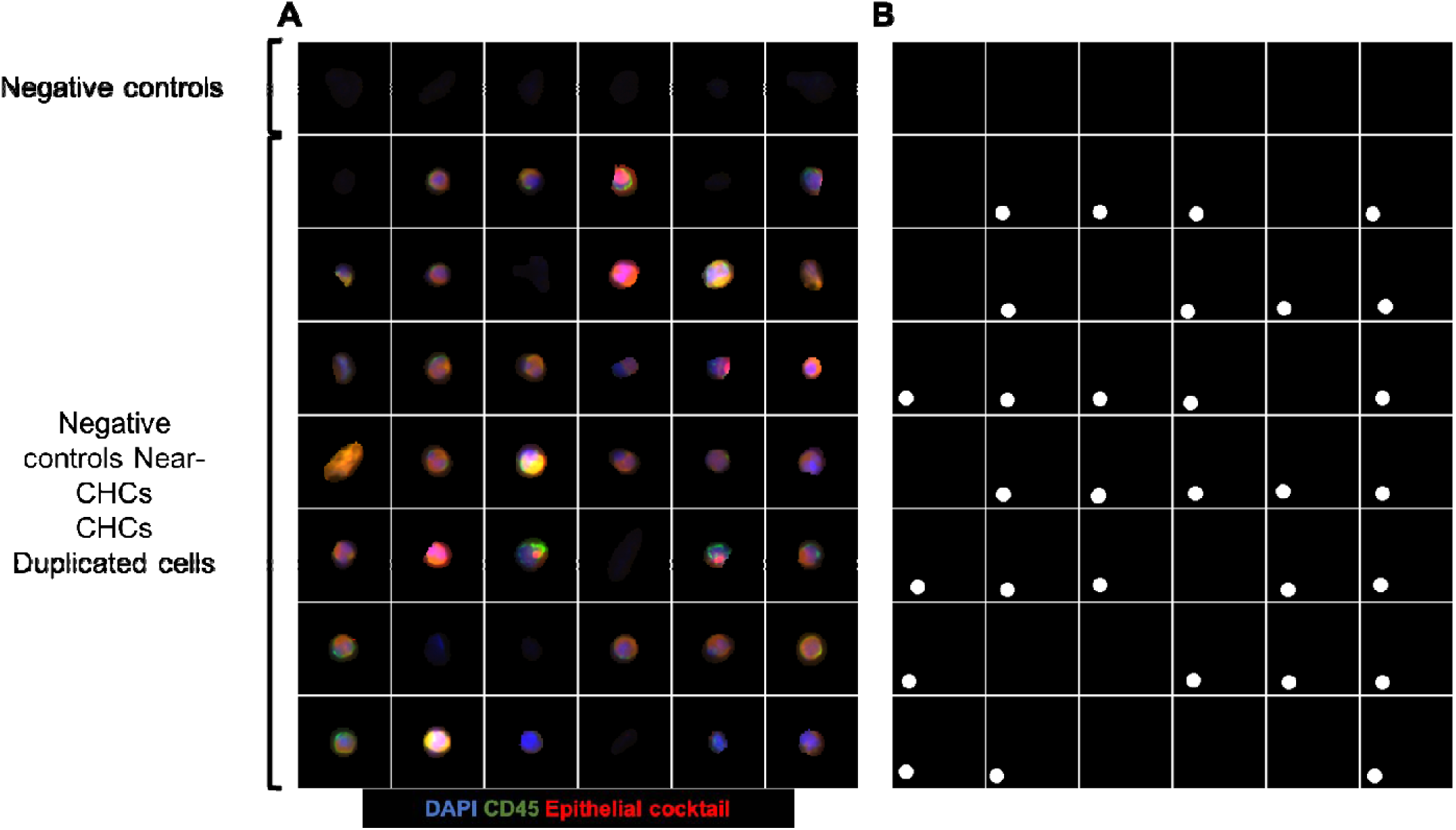
Annotation study mosaic image. Example mosaic image for 1 ROI **(A)** with labeled negative controls and corresponding annotations **(B)** created using ImageJ software.

**Figure S7:**
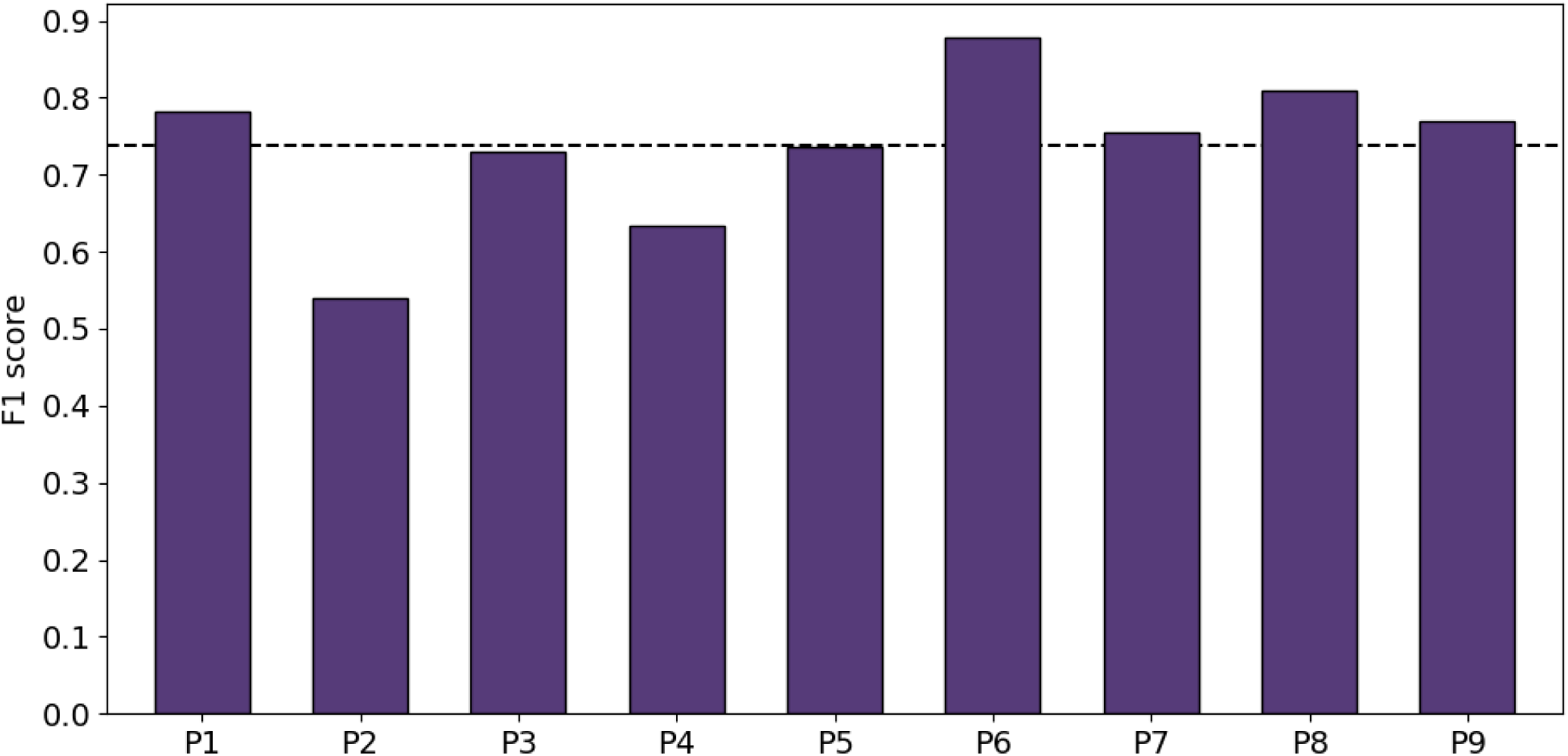
F1 scores of the 9-fold patient holdout cross validation. The dashed line represents the average F1 score across all held-out patients.

**Figure S8:**
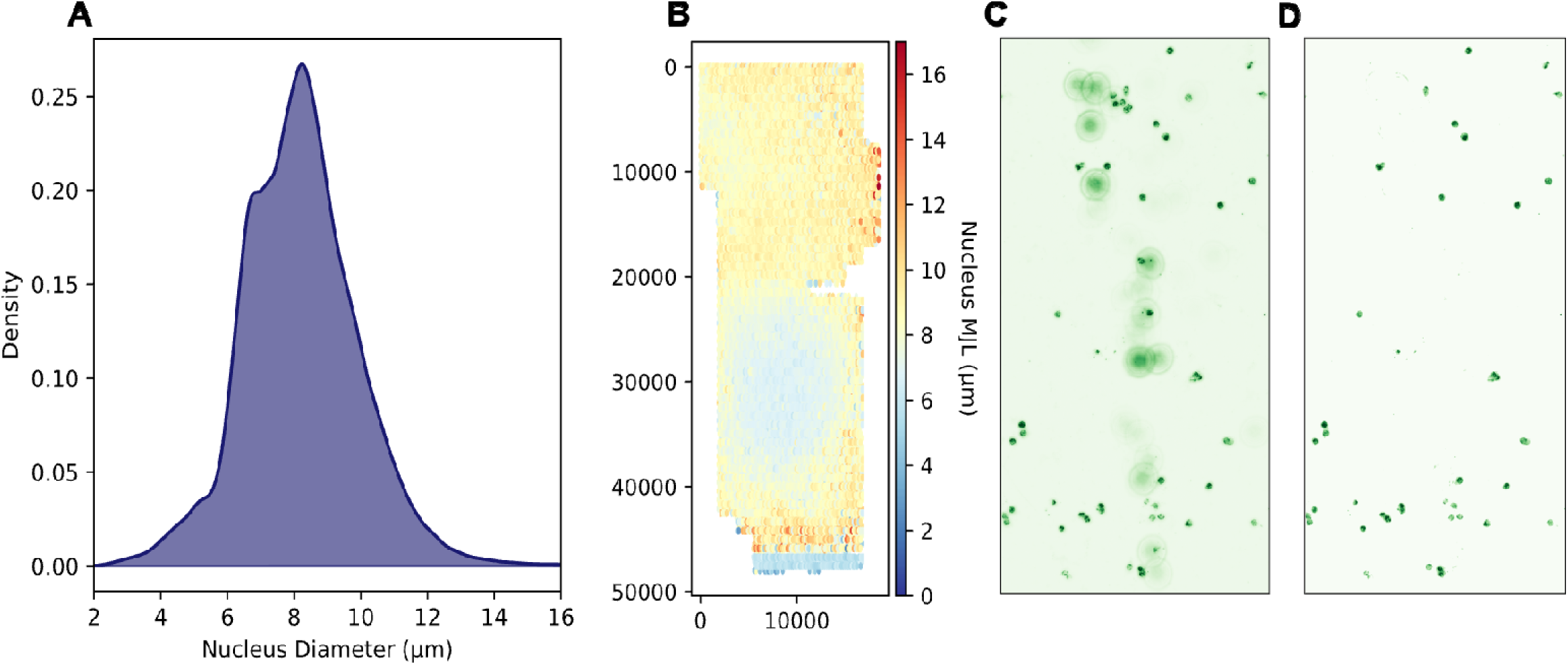
Quality control overview. **(A)** Irregular major axis length (nucleus diameter) distribution. **(B)** QC heatmaps showing uneven spatial distribution of nucleus major axis length (*um*) for the ROI shown in (A). **(C)** CD45 channel before and **(D)** after IF contaminant removal.

**Figure S9:**
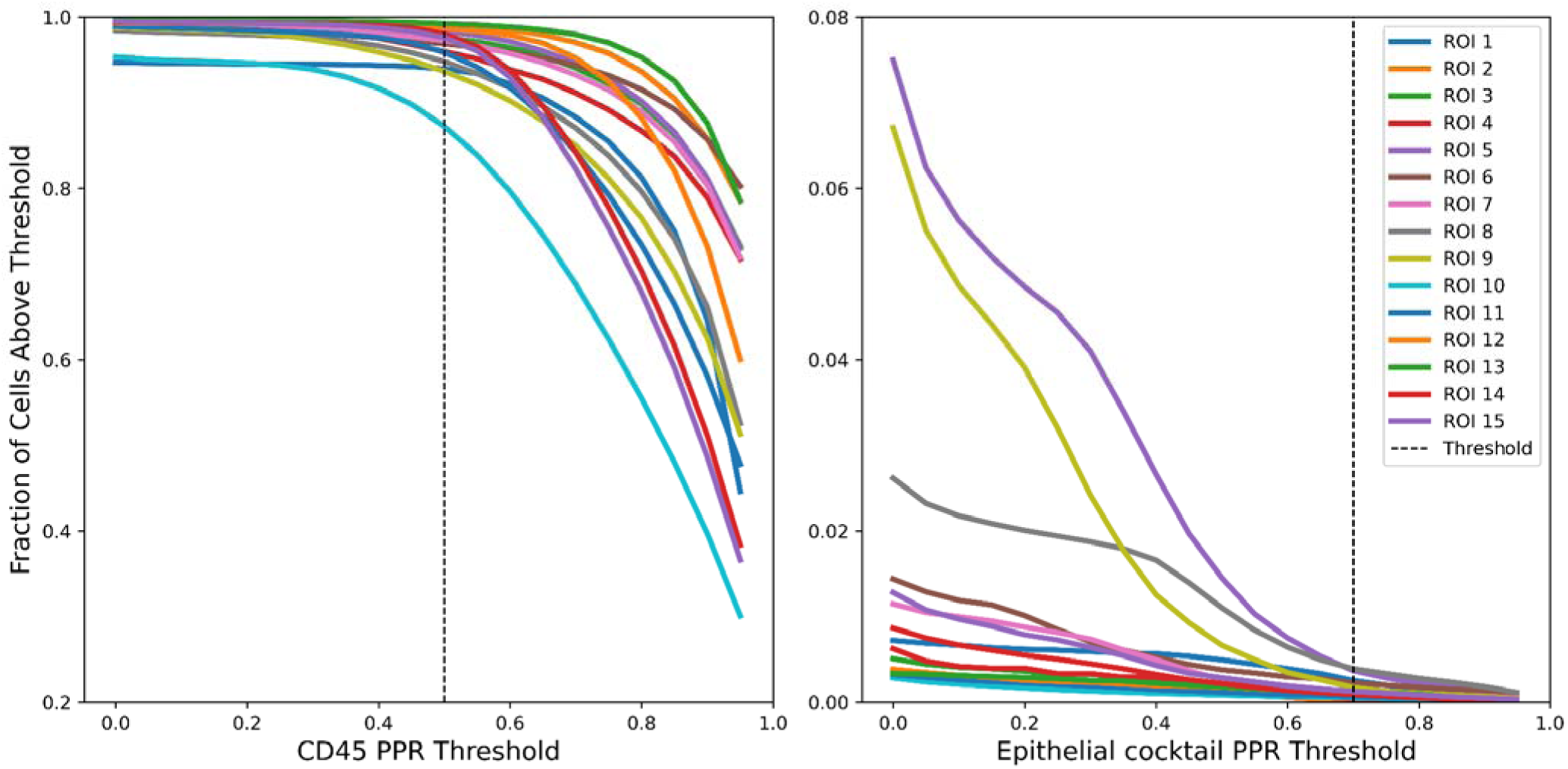
Positive pixel ratio (PPR) elbow plots. PPR elbow plots for **(A)** CD45 and **(B)** epithelial cocktail.

**Figure S10:**
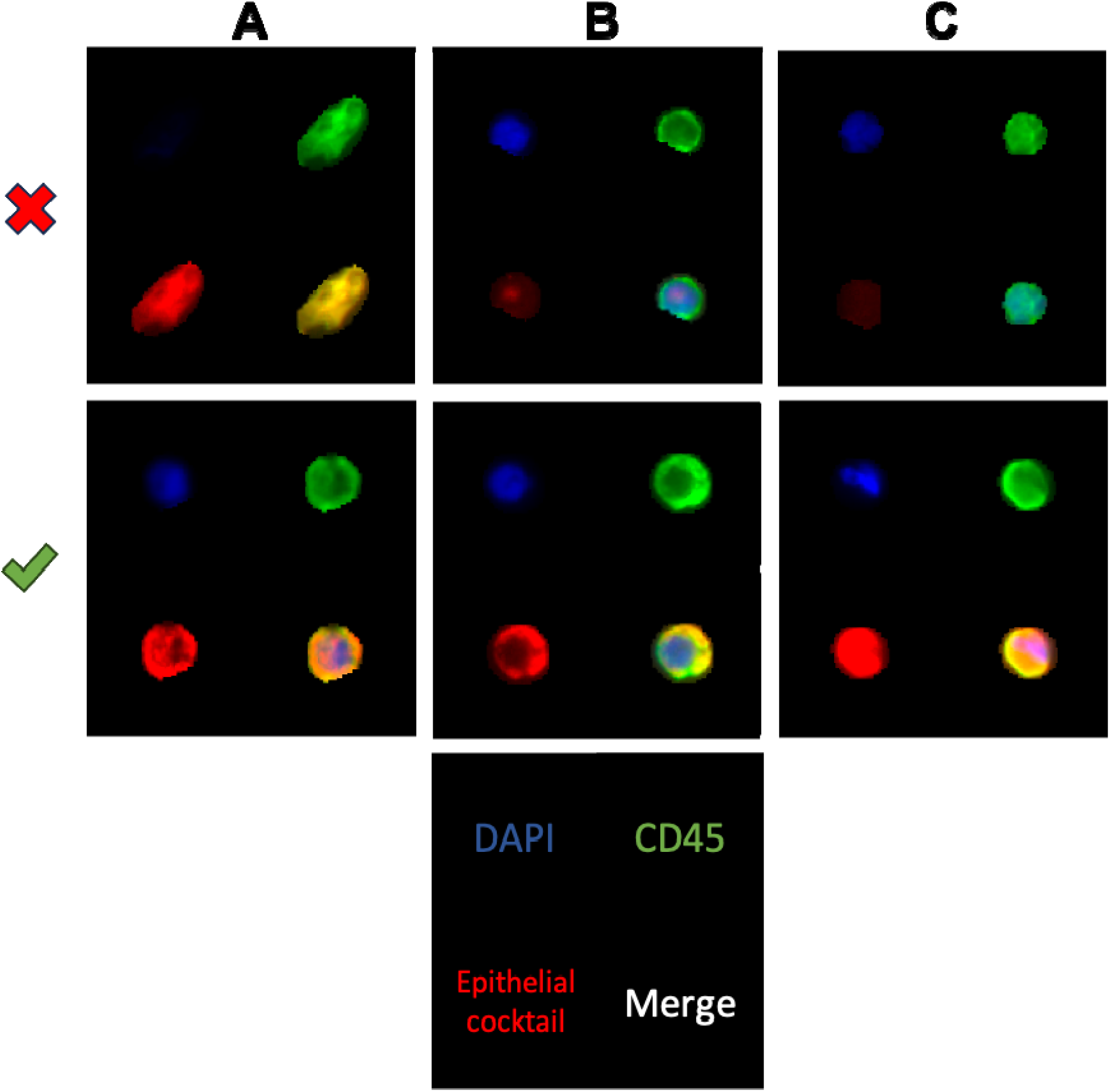
Annotator criterion for CHCs. Examples of cells that pass and fail the annotator’s criterion for calling CHCs. **(A)** DAPI intensity, **(B)** Epithelial cocktail staining pattern, and **(C)** Epithelial cocktail staining intensity.

